# Assessing wheat growth promotion potential of *Delftia lacustris* strain NSC through genomic and physiological characterization

**DOI:** 10.1101/2025.02.26.640487

**Authors:** Pinki Sharma, Rajesh Pandey, Nar Singh Chauhan

**Affiliations:** Department of Biochemistry, Maharshi Dayanand University, Rohtak, Haryana, India; INtegrative GENomics of HOst-PathogEn (INGEN-HOPE) laboratory, CSIR-Institute of Genomics and Integrative Biology (CSIR-IGIB), Mall Road, Delhi-110007, India; Academy of Scientific and Innovative Research (AcSIR), Ghaziabad-201002, India

**Keywords:** *Delftia*, Rhizosphere Microbiota, Sustainable agriculture, Biofertilizer, Biocontrol agents, Wheat Yield, Food Security

## Abstract

**Background:** *Delftia* sp. has gained considerable attention for its biocontrol and biofertilizer potential to promote the growth of crops such as *Oryza sativa*, *Brassica campestris,* and *Solanum lycopersicum*. However, *Delftia* sp. supporting wheat plants is yet to be explored.

**Methods:** *Delftia* strain was cultured from wheat rhizosphere using different growth conditions. The biofertilizer potential of *Delftia* strain was assessed through physiological, biochemical, genomic, and field experiments.

**Results and Discussion:** Phylogenetic and phylogenomic analysis confirmed taxonomic affiliation with *Delftia lacustris. In vitro* phosphate solubilization (0.325IU), nitrate reduction (0.401 IU), IAA production (0.485 IU), ACC deaminase activity (0.512 EU), siderophore synthesis properties, and strong antifungal activity against *Fusarium oxysporum* and *Rhizoctonia solani* indicated its potential as an effective biofertilizer and biocontrol agent. Drought stress tolerance (up to 40% PEG), metal stress tolerance, salt stress tolerance (up to 11.69% NaCl (w/v), 11.18% KCl (w/v), 4.24% LiCl (w/v)) indicated its survivability even in hostile conditions. Genes for phosphate solubilization (PhoR, PhoB, PhoU, PstABCD), nitrogen fixation (nifC, nifU), auxin production, siderophore biosynthesis, rhizosphere colonization and antifungal properties (chitinase, PhnZ) explains *Delftia lacustris* bioactivities. Significantly enhanced seed germination (93.33% ± 0.23), seedling growth, and biomass (*P <0.05*), particularly under stress conditions, indicated plant growth promotion properties of the *Delftia* strain. Significantly improved plant growth and yield parameters (*P=0.0001*) in field experiments, highlighting its potential as a biofertilizer and biocontrol agent.

**Conclusion:** *Delftia lacustris* strain NSC1 exhibits multifaceted biofertilizer and biocontrol potential, promoting plant growth, suppressing pathogens, and enhancing stress resilience. Its eco-friendly properties and field efficacy make it a promising alternative to chemical fertilizers for sustainable wheat production.

## Introduction

*Delftia* species are motile, aerobic, non-fermentative, gram-negative rods that belong to the Betaproteobacteria of the phylum Proteobacteria (Shigematsu et al., 2003). These microorganisms have been isolated from a variety of environments, including polluted soils, wastewater, plant rhizosphere, plant endospheres, and clinical samples, reflecting their broad metabolic versatility and saprophytic nature. *Delftia* sp. possesses nitrogen-fixing capabilities (Agafonova et al., 2017), acting as free-living diazotrophs (Han et al., 2005), while others are capable of colonizing plant roots in a symbiotic relationship (Brambilla et al., 2022), which further enhances their role in plant growth promotion. Consequently, *Delftia* sp. has attracted considerable interest in both plant growth promotion and bioremediation fields (Ubalde et al., 2012). *Delftia deserti*, *Delftia litopenaei*, *Delftia deserti*, *Delftia lacustris*, have been utilized as agricultural inoculants, bioremediation agents, and for the mineralization of toxic metals (Braña et al., 2016). *Delftia lacustris* MS3 has been isolated from heavy metal-contaminated environments, such as soils contaminated with lead or chromium (Samimi and Moghadam, 2021). *Delftia tsuruhatensis* was found to improve rice growth, particularly under nitrogen-limiting conditions, by aiding in the bioavailability of nutrients and enhancing root development, thus contributing to overall plant health and productivity (Harahap et al., 2023). *Paraburkholderia fungorum* BRRh-4 and *Delftia* sp. BTL-M2 has been shown to promote plant growth and increase yield (Islam et al., 2023) in *Oryza sativa*. These strains have demonstrated the ability to produce siderophores (Guo et al., 2016), which aid in the chelation and mobilization of metal ions and have been shown to promote plant growth, thereby highlighting the potential for these microorganisms to simultaneously contribute to both bioremediation and plant health improvement (Samimi and Shahriari-Moghadam, 2021).

Despite their well-documented role in enhancing plant growth, improving stress resilience, and supporting nutrient uptake, the application of *Delftia* strains in wheat cultivation has not been extensively studied. Wheat rhizosphere was identified as a potential source of biofertilizer and biocontrol strains. *Delftia* strain from wheat rhizosphere could exhibit biofertilizer and biocontrol properties and be successfully employed to enhance wheat growth. Hereby, the present study aimed to isolate *Delftia* strain with a broader, integrated capacity to promote plant growth while concurrently providing protection against both biotic and abiotic stressors. The focus is on identifying *Delftia* strain that supports nutrient uptake and plant development and enhances the plant’s resilience to environmental stresses such as drought, salinity, and pathogen attacks, thus offering a holistic approach to improving plant health and productivity.

## Material and Methods

### Isolation of wheat rhizosphere microbes (WRMs)

Wheat rhizospheric microbes were isolated using previously standardized methods (Sharma et al., 2024a). Phylogenetic affiliation of the isolated microbial strains was assessed through 16S rRNA gene-based phylotype identification (Sharma et al., 2024b)

### Molecular, physiological, biochemical, and genomic characterization of *Delftia* strain NSC

Gram staining was performed with a gram staining kit (K001-1KT, Himedia). The growth was observed at different pH ranges (3, 4, 5, 7, 8, 9, 10, 11, and 12) and temperature ranges (10°C, 20°C, 30°C, 40°C, 50°C, 60°C) to identify its optimal growth conditions. Its growth pattern was observed in LB broth for 48 hours at 37°C with constant shaking at 200 to check its doubling time (Sharma et al., 2024a). Substrate utilization preference was assessed with a Hi carbo kit (Himedia, KB009A-1KT, KB009B-1KT, and KB009C-1KT). Biochemical properties were performed using assays for amylase (Swain et al., 2006), catalase (Iwase et al., 2013), pectinase (Oumer and Abate, 2018), cellulase (Kasana et al., 2008), esterase (Ramnath et al., 2017), and protease Vijayaraghavan et al., 2017). The stress response physiology was assessed by performing specific assays for salt stress tolerance, metal stress tolerance, and oxidative stress (Sharma et al., 2024). DNA was extracted from the microbial isolate using the alkali lysis method (Chauhan et al., 2009). The qualitative and quantitative analysis of the DNA was performed with agarose gel electrophoresis and Qubit HS DNA estimation kits (Invitrogen, USA), respectively. Genomic DNA was sequenced using Illumina MiSeq platform using Nextera XT DNA Library Prep kit (Yadav et al., 2022) to unveil its genomic architecture. Quality filtration, genome assembly, and genome completeness were checked using standard methodology (Yadav et al., 2022). Genomic characterization and comparative genomics were performed against other *Delftia* using a standardized methodology (Yadav et al., 2022).

### Assessment of Biofertilizer-related features in *Delftia lacustris strain* NSC

*Delftia lacustris* strain NSC was screened for phosphate solubilization (Behera et al., 2017), nitrate reductase activity (Kim and Seo, 2018), Indole-3-acetic acid (IAA) production (Ehmann, 1977), ammonia production (Bhattacharyya et al., 2020), and siderophore biosynthesis (Himpsl and Mobley, 2019) to assess its bio-fertilization potential. Drought stress tolerance was evaluated as described by Elizabeth Mustamu et al. (2023) (Elizabeth Mustamu et al., 2023). The salt-induced oxidative stress was evaluated by measuring ACC deaminase production activity, following a standardized methodology (Maheshwari et al., 2020). The RAST software assessed the genetic features related to plant growth promotion (https://rast.nmpdr.org/rast.cgi).

### Assessment of *Delftia lacustris strain* NSC biofertilizer and biocontrol properties in laboratory conditions

The biocontrol potential was evaluated against *Rhizoctonia solani,* and *Fusarium oxysporum*. The assessment focused on their effects on seed germination efficiency and wheat seedlings’ root and shoot lengths (Sharma et al., 2024a; Singh and Kayastha, 2014). To investigate the role of *Delftia lacustris* strain NSC in seed germination under saline conditions, seeds were soaked in an overnight-grown microbial culture (cell density of 10^11^ cells/mL), supplemented with NaCl concentrations ranging from 0 to 1 M, for 16 hours at 37°C. Control seeds were soaked directly in NaCl solutions of equivalent concentrations for 16 hours at 37°C. Following the soaking treatment, seeds were wrapped in germination sheets and placed in 50 mL culture tubes containing 5 mL of Hoagland solution. The tubes were incubated in the dark at ambient temperature (25°C) for 7 days. After incubation, wheat seed germination percentage, alpha-amylase activity, and root and shoot lengths were measured (Sharma et al., 2024a; Singh and Kayastha, 2014).

### Assessment of *Delftia lacustris strain* NSC biofertilizer properties under field conditions

Roots of WC-306 plants treated with *Delftia lacustris* strain NSC were harvested at different Feeks and analyzed for their phosphate-solubilizing ability (Behera et al., 2017), nitrate reductase activity (Kim and Seo, 2018), total sugar content (Ludwig and Goldberg, 1956), and reducing sugar content (Khatri and Chhetri, 2020), with comparisons made to untreated WC-306 plants. Additionally, various growth and yield parameters were assessed in both treated and untreated plants, including seed germination percentage, the number of tillers per plant, number of leaves per plant, number of spikes per plant, spike length, number of spikelets per spike, and grain yield, to evaluate the effects of *Delftia lacustris* strain NSC on plant growth and productivity.

### Statistical analysis

All experiments were performed with replicate samples. Statistical analyses and data visualization were conducted using SIGMA Plot 15. Analysis of variance (ANOVA) was applied to determine the significance between the microbial-treated and untreated groups [Systat Software SigmaPlot 15].

## Results

### Identification of microbial isolate NSC

Microbe was isolated using the standard methodology described in our previous study (Sharma et al., 2024). The microbial isolates were obtained from the wheat rhizosphere having a pH of 7.3 ± 0.0025, a temperature of 22.6°C ± 0.01, and a moisture content of 11.5 ± 1.10014. The 16S rRNA gene-based taxonomic characterization revealed that the NSC isolatès 16S rRNA gene shared 99.86% homology with *Delftia lacustris* strain 332 (NR 116495.1) in the NCBI Nr database. This identification was further confirmed through phylogenetic analysis (**Figure 1A**).

**Figure 1:**
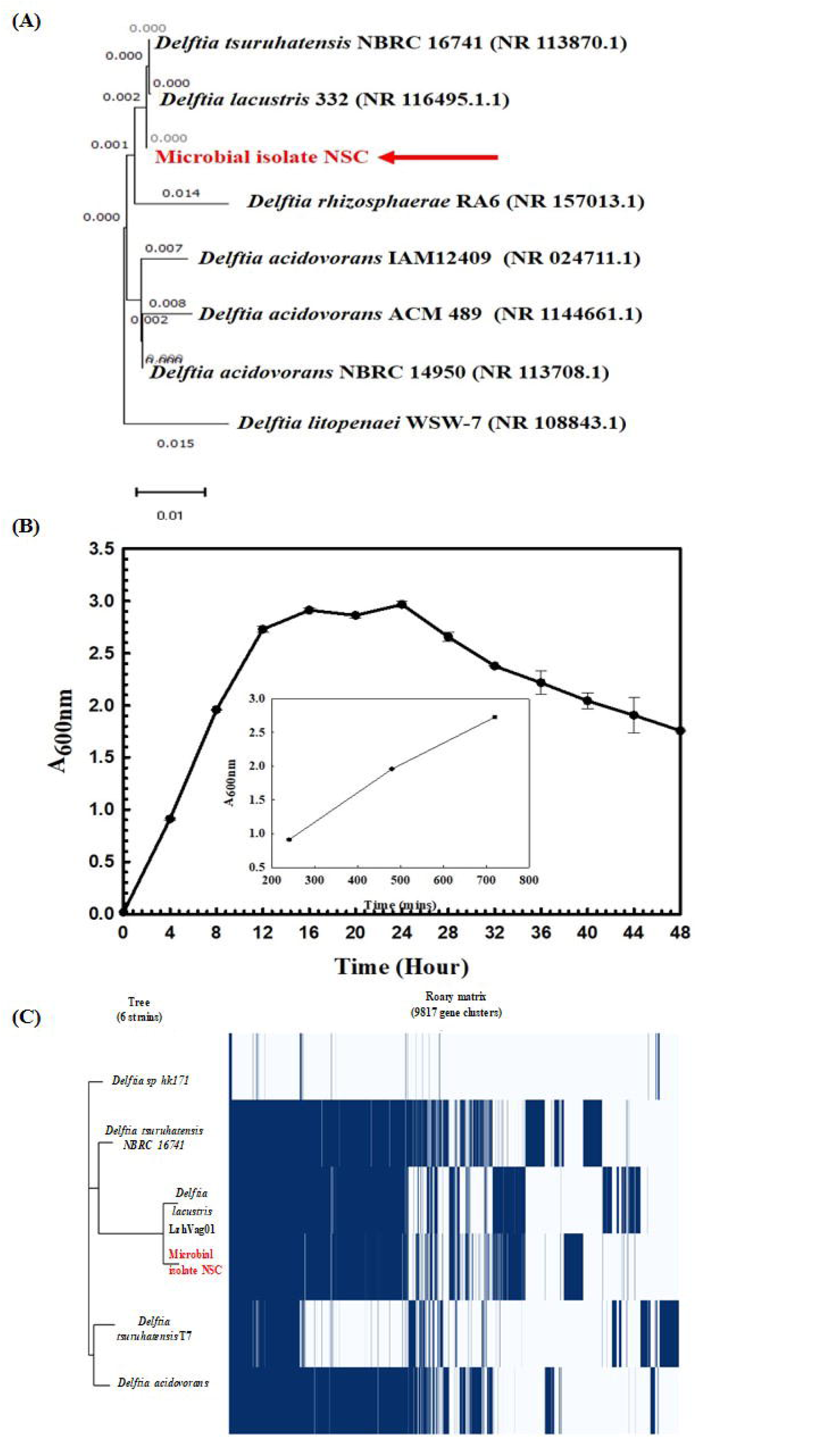
Phylogenetic and Physiological characterization of *Delftia lacustris* strain NSC. Phylogenetic affiliation of microbial isolates NSC with the other *Delftia* species (A). The phylogenetic tree was constructed using the Neighbor-joining method of phylogenetics with 1000 bootstrap replications using MEGA-X software. The out-group was represented by *Delftia litopenaei* WSW-7 SSU rRNA gene sequence. Growth pattern analysis of *Delftia lacustris* strain NSC was observed after incubating the cultures for 48 hours in LB broth with constant shaking at 200 rpm (B). Phylogenomic analysis of *Delftia lacustris* strain NSC with other *Delftia* strains (C). The phylogenomic tree of the microbial isolate was generated using the FastTree v2.1.10 tool through Roary. The left panel illustrates the phylogenetic relationship of *Delftia lacustris* NSC with other *Delftia* species, while the right panel shows the core and accessory genes shared among different *Delftia lacustris* strains

### Molecular, physiological, biochemical, and genomic characterization of *Delftia* strain NSC

Microscopic examination identified the isolate as a gram-negative, rod-shaped, motile bacterium. Physiologically, it exhibited optimal growth at pH 7.0 and 35°C temperature, reaching the logarithmic phase after 12 hours with a doubling time of approximately 54.31 minutes (**Figure 1B**). It also demonstrated facultative anaerobic growth, with an optical density (O.D.) of 0.402 at 600 nm after 24 hours under anaerobic conditions. Biochemical testing showed it was positive for amylase, esterase, lipase, protease, and catalase activities, and its substrate utilization profile was consistent with that of other *Delftia* species (**Supplementary Table S1**). Antibiotic susceptibility testing revealed that its resistance to amoxicillin, bacitracin, cephalothin, vancomycin, ceftazidime, and ofloxacin, but remained sensitive to oxytetracycline, novobiocin, and lincomycin, resembling the antibiotic resistance profile of other *Delftia* strains (**Supplementary Table S2**). Stress response assays demonstrated that it could tolerate saltine stress (up to 11.69% NaCl, 11.18% KCl, 4.24% LiCl), metal stress (0.068% CdCl_2_, up to 0.1% Na_3_AsO_4_, 0.15% NaAsO_2_), and oxidizing agents stress (up to 4.25% H_2_O_2_), similar to other *Delftia* strains (**Supplementary Table S3**). These findings highlight the adaptability of *Delftia* strain NSC under stress conditions. Genome sequencing *Delftia lacustris* strain NSC generated 805,929 paired-end raw reads, which were assesmbled 6,235,469 base pairs represented through 220 contigs. Functional annotation of genome revealed 5,466 protein-coding sequences, 8 rRNA genes, 82 tRNA genes, and one tmRNA gene. The Average Nucleotide Identity (ANI) among different species of *Delftia* sp. ranged from 74% to 98%, reflecting significant genomic variation between species. Notably, the ANI between *Delftia lacustris* NSC and *Delftia lacustris* LzhVag01 was 98.28%, higher than those observed with other species within the genus (**Supplementary Table S4**). The classification of *Delftia lacustris* strain NSC as part of the *Delftia* species was further confirmed using terra correlation. *Delftia* sp. LMG 24775 showed a z-score of 0.99989 when compared to *Delftia lacustris* strain NSC, confirming its close genetic relationship. Other *Delftia* species exhibited high similarity, with z-scores ranging from approximately 0.95 to 0.99 (**Supplementary Table S5**). Following ANIb and tetra confirmation, the genome of *Delftia lacustris* strain NSC was compared to those of other *Delftia* species to assess genome-wide similarities and differences. A matrix generated using the Roary tool highlighted the extensive nature of the genome, showing that *Delftia lacustris* strain NSC exhibited the greatest similarity to *Delftia lacustris* LzhVag01 (**Figure 1C**). The Roary analysis also revealed that all *Delftia* genomes shared a significant number of genes. The core genome, along with the shell and cloud genomes, form the fundamental genetic structure of these organisms, with the core genes providing essential functions, while the additional genomes contribute to variability and adaptability to various environmental conditions or ecological niches. Notably, the *Delftia lacustris* strain NSC genome lacks pathogenic genes or virulence-related genes, supporting its non-pathogenic nature. Genomic analysis also identified genes involved in phosphate solubilization and transport (**Table 1**). Beyond phosphate solubilization, its genome includes genes associated with plant growth promotion activities, such as auxin biosynthesis, nitrogen assimilation, and siderophore biosynthesis (**Table 1**). Additionally, genes for arsenic resistance, oxidative stress tolerance, metal stress tolerance, and salt tolerance were present, highlighting its capacity for stress response. Several proteins essential for effective colonization of the plant rhizosphere were identified (Kumar et al., 2023). A detailed examination of the *Delftia lacustris* strain NSC genomes also revealed genes encoding proteins for the synthesis of Type 1 & IV pili and exopolysaccharides (**Table 1**), crucial for plant surface adhesion, auto-aggregation, and early biofilm formation (Sharma et al., 2024).

**Table 1:**
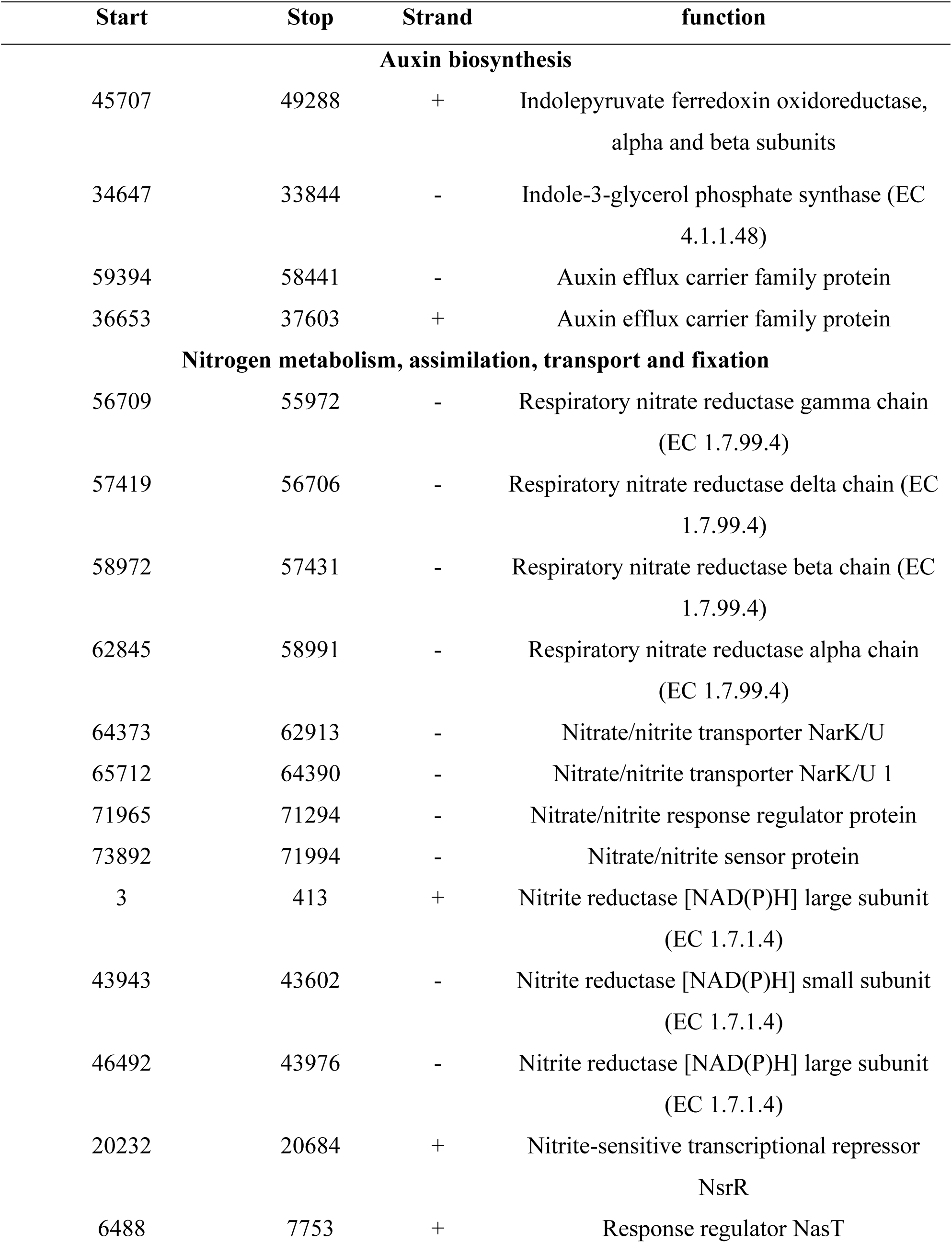
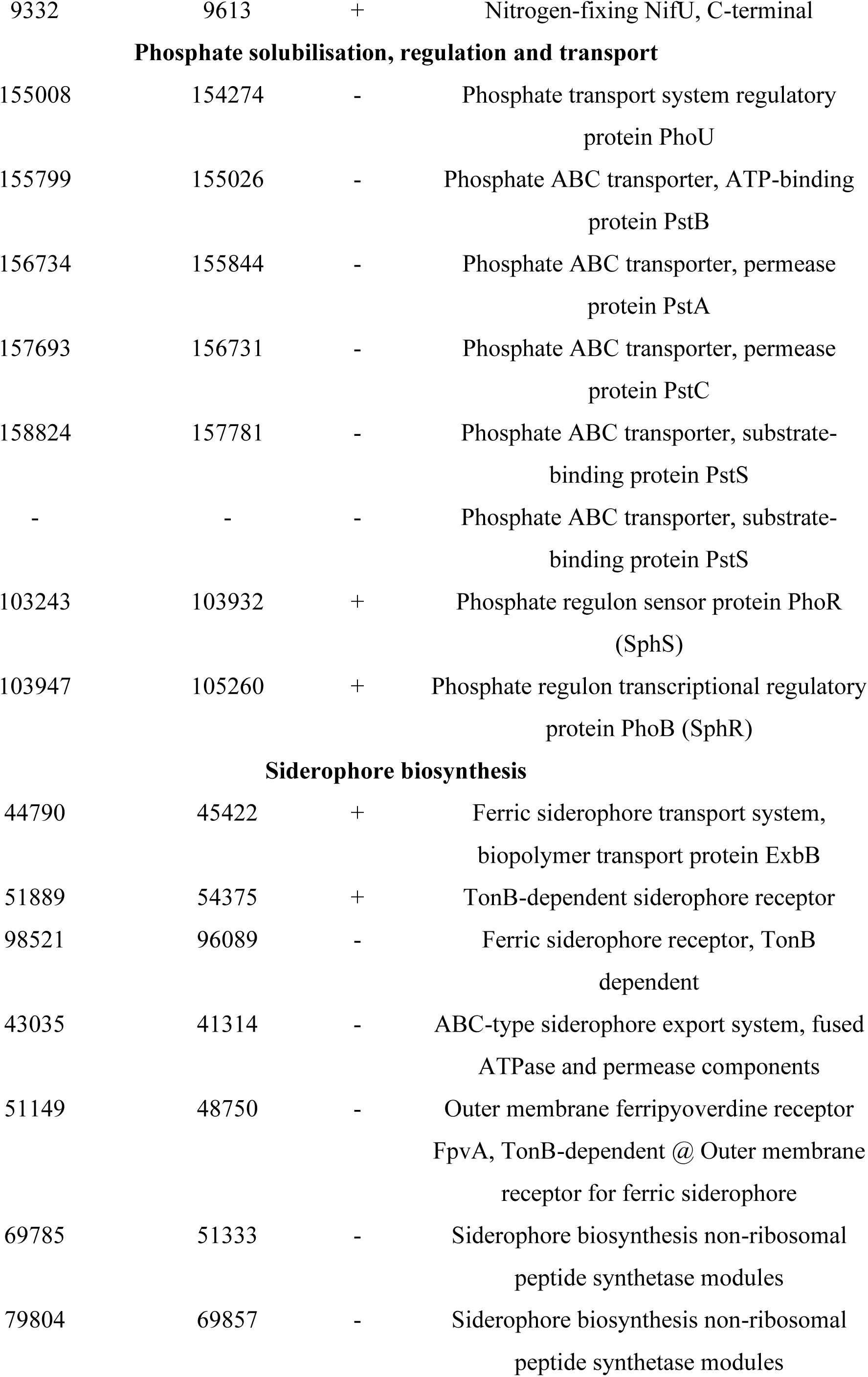
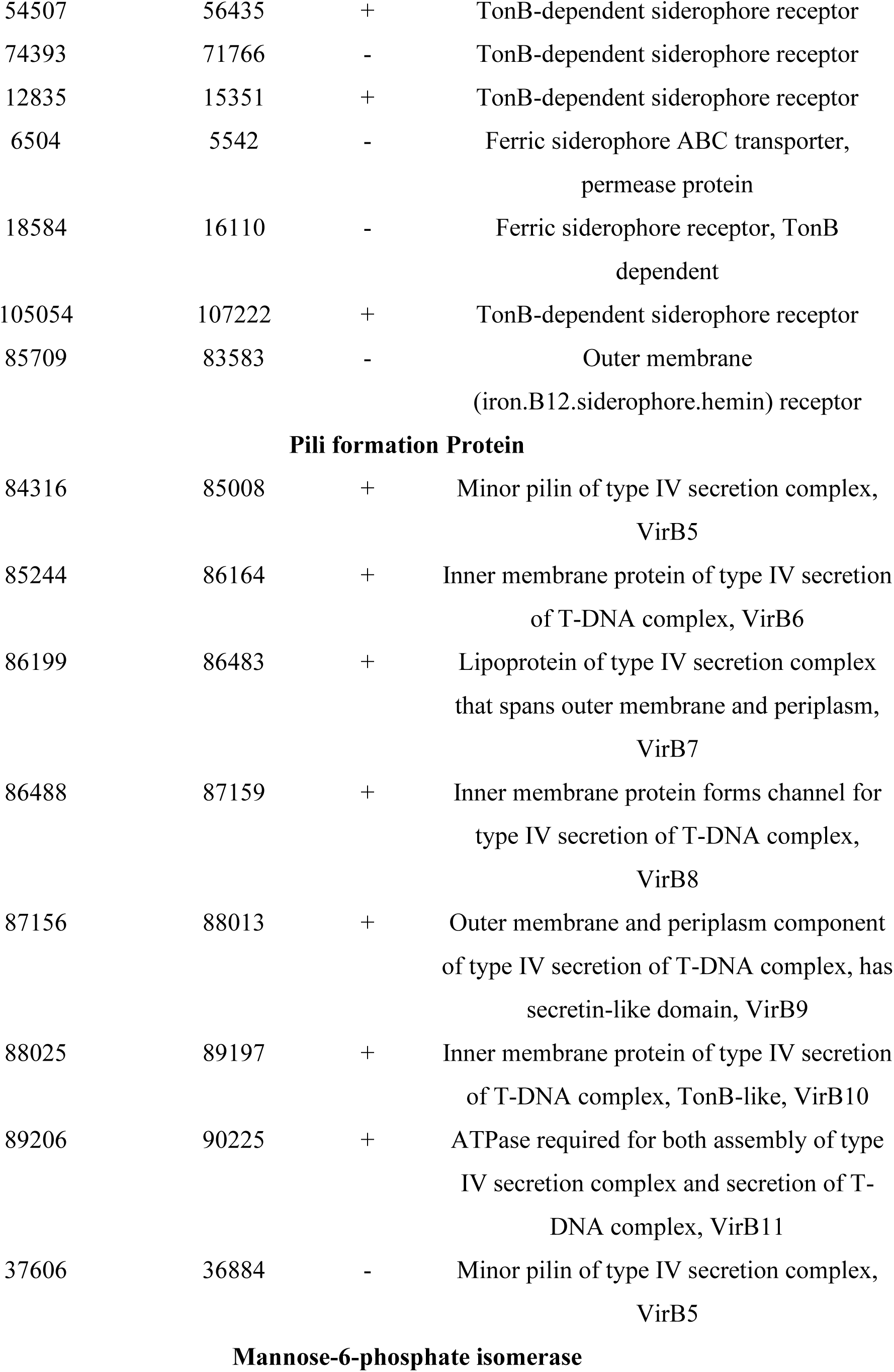
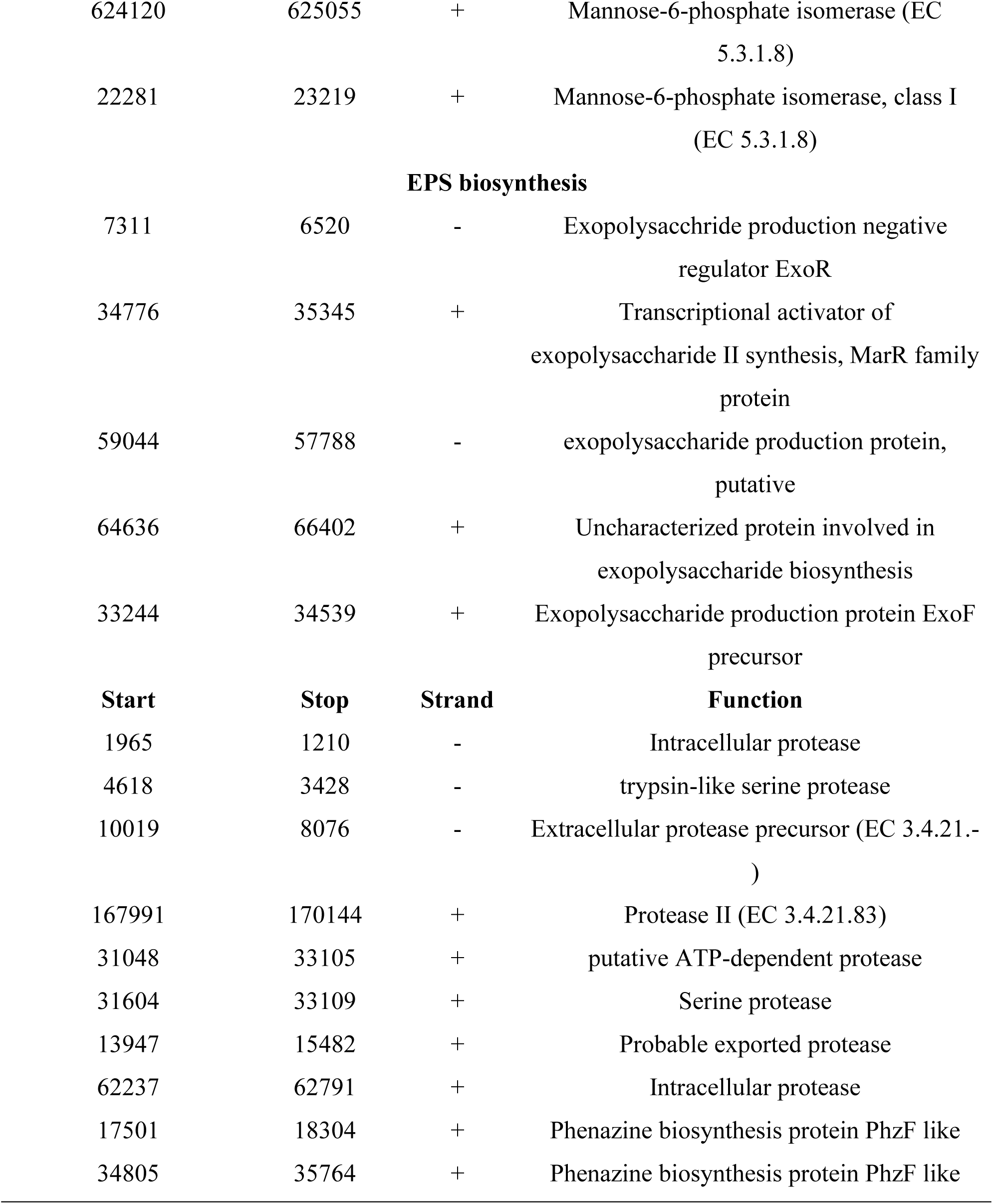
Genetic features identified within *Delftia lacustris* strain NSC genome encoding various proteins involved in nutrient assimilation, solubilization, colonization, and antifungal properties.

| Start | Stop | Strand | function |
| --- | --- | --- | --- |
| **Auxin biosynthesis** |  |  |  |
| 45707 | 49288 | + | Indolepyruvate ferredoxin oxidoreductase, alpha and beta subunits |
| 34647 | 33844 | - | Indole-3-glycerol phosphate synthase (EC 4.1.1.48) |
| 59394 | 58441 | - | Auxin efflux carrier family protein |
| 36653 | 37603 | + | Auxin efflux carrier family protein |
| **Nitrogen metabolism, assimilation, transport and fixation** |  |  |  |
| 56709 | 55972 | - | Respiratory nitrate reductase gamma chain (EC 1.7.99.4) |
| 57419 | 56706 | - | Respiratory nitrate reductase delta chain (EC 1.7.99.4) |
| 58972 | 57431 | - | Respiratory nitrate reductase beta chain (EC 1.7.99.4) |
| 62845 | 58991 | - | Respiratory nitrate reductase alpha chain (EC 1.7.99.4) |
| 64373 | 62913 | - | Nitrate/nitrite transporter NarK/U |
| 65712 | 64390 | - | Nitrate/nitrite transporter NarK/U 1 |
| 71965 | 71294 | - | Nitrate/nitrite response regulator protein |
| 73892 | 71994 | - | Nitrate/nitrite sensor protein |
| 3 | 413 | + | Nitrite reductase [NAD(P)H] large subunit (EC 1.7.1.4) |
| 43943 | 43602 | - | Nitrite reductase [NAD(P)H] small subunit (EC 1.7.1.4) |
| 46492 | 43976 | - | Nitrite reductase [NAD(P)H] large subunit (EC 1.7.1.4) |
| 20232 | 20684 | + | Nitrite-sensitive transcriptional repressor NsrR |
| 6488 | 7753 | + | Response regulator NasT |
| 9332 | 9613 | + | Nitrogen-fixing NifU, C-terminal |
| **Phosphate solubilisation, regulation and transport** |  |  |  |
| 155008 | 154274 | - | Phosphate transport system regulatory protein PhoU |
| 155799 | 155026 | - | Phosphate ABC transporter, ATP-binding protein PstB |
| 156734 | 155844 | - | Phosphate ABC transporter, permease protein PstA |
| 157693 | 156731 | - | Phosphate ABC transporter, permease protein PstC |
| 158824 | 157781 | - | Phosphate ABC transporter, substrate-binding protein PstS |
| - | - | - | Phosphate ABC transporter, substrate-binding protein PstS |
| 103243 | 103932 | + | Phosphate regulon sensor protein PhoR (SphS) |
| 103947 | 105260 | + | Phosphate regulon transcriptional regulatory protein PhoB (SphR) |
| **Siderophore biosynthesis** |  |  |  |
| 44790 | 45422 | + | Ferric siderophore transport system, biopolymer transport protein ExbB |
| 51889 | 54375 | + | TonB-dependent siderophore receptor |
| 98521 | 96089 | - | Ferric siderophore receptor, TonB dependent |
| 43035 | 41314 | - | ABC-type siderophore export system, fused ATPase and permease components |
| 51149 | 48750 | - | Outer membrane ferripyoverdine receptor FpvA, TonB-dependent @ Outer membrane receptor for ferric siderophore |
| 69785 | 51333 | - | Siderophore biosynthesis non-ribosomal peptide synthetase modules |
| 79804 | 69857 | - | Siderophore biosynthesis non-ribosomal peptide synthetase modules |
| 54507 | 56435 | + | TonB-dependent siderophore receptor |
| 74393 | 71766 | - | TonB-dependent siderophore receptor |
| 12835 | 15351 | + | TonB-dependent siderophore receptor |
| 6504 | 5542 | - | Ferric siderophore ABC transporter,<br>permease protein |
| 18584 | 16110 | - | Ferric siderophore receptor, TonB<br>dependent |
| 105054 | 107222 | + | TonB-dependent siderophore receptor |
| 85709 | 83583 | - | Outer membrane<br>(iron.B12.siderophore.hemin) receptor |
| **Pili formation Protein** |  |  |  |
| 84316 | 85008 | + | Minor pilin of type IV secretion complex,<br>VirB5 |
| 85244 | 86164 | + | Inner membrane protein of type IV secretion<br>of T-DNA complex, VirB6 |
| 86199 | 86483 | + | Lipoprotein of type IV secretion complex<br>that spans outer membrane and periplasm,<br>VirB7 |
| 86488 | 87159 | + | Inner membrane protein forms channel for<br>type IV secretion of T-DNA complex,<br>VirB8 |
| 87156 | 88013 | + | Outer membrane and periplasm component<br>of type IV secretion of T-DNA complex, has<br>secretin-like domain, VirB9 |
| 88025 | 89197 | + | Inner membrane protein of type IV secretion<br>of T-DNA complex, TonB-like, VirB10 |
| 89206 | 90225 | + | ATPase required for both assembly of type<br>IV secretion complex and secretion of T-DNA complex, VirB11 |
| 37606 | 36884 | - | Minor pilin of type IV secretion complex,<br>VirB5 |
| **Mannose-6-phosphate isomerase** |  |  |  |
| 624120 | 625055 | + | Mannose-6-phosphate isomerase (EC 5.3.1.8) |
| 22281 | 23219 | + | Mannose-6-phosphate isomerase, class I (EC 5.3.1.8) |
| **EPS biosynthesis** |  |  |  |
| 7311 | 6520 | - | Exopolysacchride production negative regulator ExoR |
| 34776 | 35345 | + | Transcriptional activator of exopolysaccharide II synthesis, MarR family protein |
| 59044 | 57788 | - | exopolysaccharide production protein, putative |
| 64636 | 66402 | + | Uncharacterized protein involved in exopolysaccharide biosynthesis |
| 33244 | 34539 | + | Exopolysaccharide production protein ExoF precursor |
| **Start** | **Stop** | **Strand** | **Function** |
| 1965 | 1210 | - | Intracellular protease |
| 4618 | 3428 | - | trypsin-like serine protease |
| 10019 | 8076 | - | Extracellular protease precursor (EC 3.4.21.- ) |
| 167991 | 170144 | + | Protease II (EC 3.4.21.83) |
| 31048 | 33105 | + | putative ATP-dependent protease |
| 31604 | 33109 | + | Serine protease |
| 13947 | 15482 | + | Probable exported protease |
| 62237 | 62791 | + | Intracellular protease |
| 17501 | 18304 | + | Phenazine biosynthesis protein PhzF like |
| 34805 | 35764 | + | Phenazine biosynthesis protein PhzF like |

### Assessment of Biofertilizer-related features in *Delftia lacustris strain* NSC

It demonstrated phosphate solubilizing activity (0.325IU), nitrate reductase activity (0.401 IU), and produced IAA (0.485 IU). The qualitative analysis of ammonia production showed that *Delftia lacustris* strain NSC was capable of generating ammonia. Siderophore biosynthesis assays revealed the formation of an orange color and the development of a clearance zone (19.5 ± 0.02 mm), indicating the siderophore production ability. Additionally, it was able to produce and secrete plant growth-promoting hormones into their environment. The presence of genes linked to plant growth promotion and their demonstrated bioactivity suggest that it has nutrient assimilation capabilities that can enhance plant growth. It has also demonstrated strong growth even in the presence of up to 40% polyethylene glycol (PEG), indicating its potential to promote plant growth under drought-stress conditions. This result suggests that *Delftia lacustris* strain NSC possesses traits that can help plants withstand drought by enhancing growth in challenging environments. Additionally, it was found to produce the enzyme ACC deaminase, with an activity level of 0.512 EU. The presence of ACC deaminase activity suggests that it may play a crucial role in promoting plant health under stressful conditions, such as drought, by reducing oxidative stress and supporting plant growth.

### Assessment of *Delftia lacustris strain* NSC biofertilizer and biocontrol properties in laboratory conditions

The protective effect of *Delftia lacustris* strain NSC on wheat seed germination was assessed. The control group demonstrated a germination rate of 75.66 ± 0.57735%. In contrast, seeds pre-treated with *Delftia lacustris* strain NSC exhibited a germination efficiency of 93.33 ± 0.23 % (**Figure 2A**), reflecting approximately a 123.35-fold increase (*P = 0.0001*) in germination percentage. In the presence of *Rhizoctonia solani* and *Fusarium oxysporum*, seed germination was substantially reduced in untreated seeds to 10 ± 1% (*P = 0.024*) and 5 ± 0.57735 % (*P = 0.032*), respectively. However, pre-treatment with *Delftia lacustris* strain NSC led to improved germination efficiencies of 52.25 ± 0.23% (*P = 0.0015*) and 48.19 ± 0.55% (*P = 0.024*) in the presence of *Rhizoctonia solani* and *Fusarium oxysporum*, respectively (**Figur**e **2A**). Notably, it has enhanced seed germination in the presence of these phytopathogens by approximately 520-fold (*P = 0.0001*) and 963.8-fold (*P = 0.0001*) compared to untreated control seeds. These findings strongly suggest the biocontrol potential of the *Delftia lacustris* strain NSC, as it not only improved seed germination in the presence of phytopathogens but also significantly outperformed untreated seeds (*P = 0.0025*) (**Figure 2**). Moreover, *Rhizoctonia solani* and *Fusarium oxysporum* significantly reduced alpha-amylase activity in wheat seeds to 0.42 IU and 0.22 IU, respectively (*P = 0.0001* and *P = 0.0018*). Pre-treatment with *Delftia lacustris* strain NSC significantly restored alpha-amylase activity to 0.792 IU (*P = 0.0001*) and 0.735 IU (*P = 0.0001*) in the presence of *Rhizoctonia solani* and *Fusarium oxysporum*, respectively (**Figure 2B**). The marked increase in alpha-amylase activity following pre-treatment with *Delftia* isolate could account for the enhanced seed germination observed.

**Figure 2:**
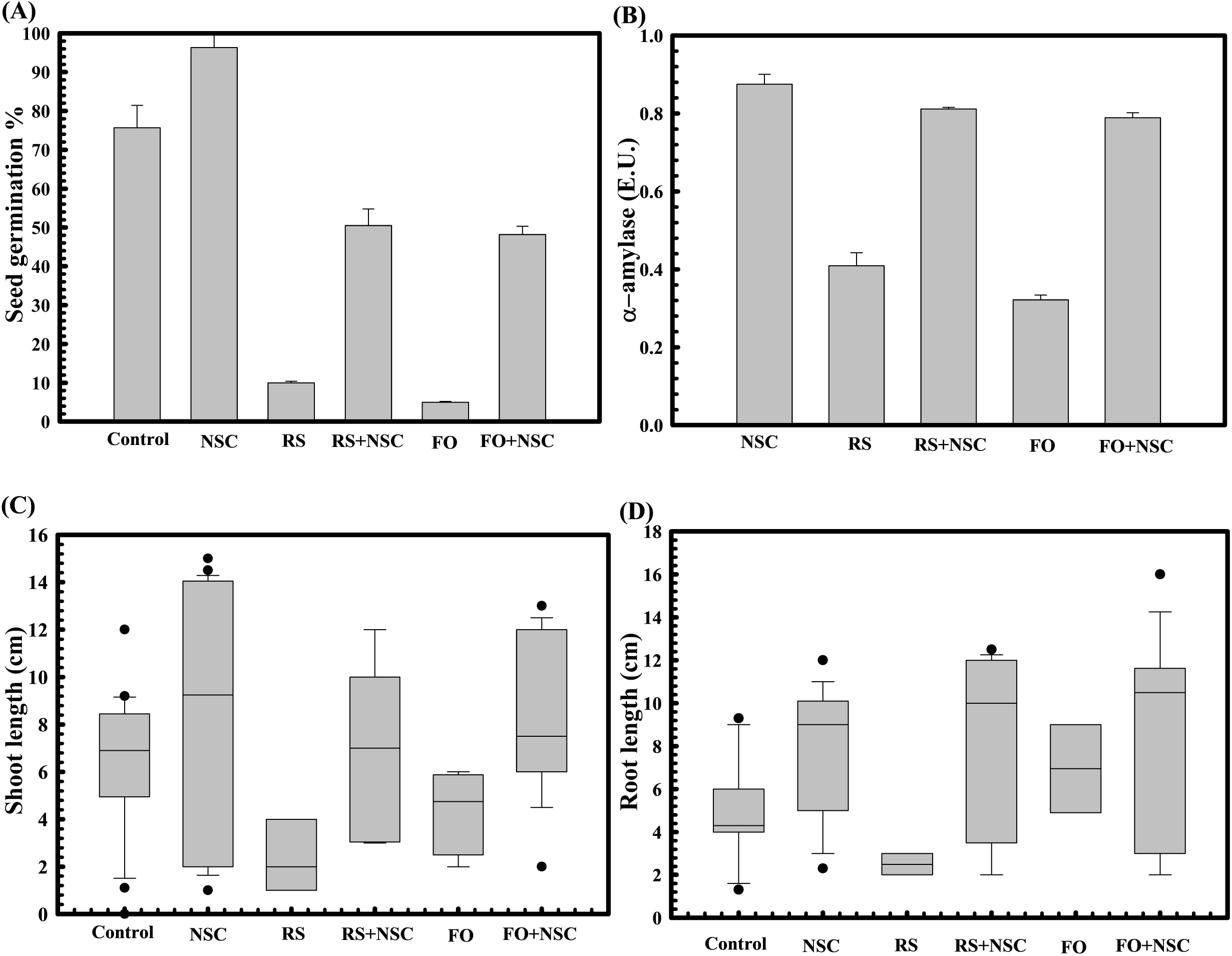
Effect of *Delftia lacustris* strain NSC on wheat seed germination parameters. The seed germination percentage in *Delftia lacustris* strain NSC pretreated seed in reference to the control (untreated) (A). The alpha amylase activity of *Delftia lacustris* strain NSC pretreated wheat seeds in the presence and absence of plant pathogenic fungi *Rhizoctonia solani* and *Fusarium oxysporum* (B). The impact of seeds pretreatment with D*elftia lacustris* strain NSC on shoot (C) and root length (D) of WC-306 seedling in the presence of *Rhizoctonia solani* and *Fusarium oxysporum*. Plotted values are the mean of triplicates along with the observed standard deviation. Here NSC: *Delftia lacustris* NSC; FO: *Fusarium oxysporum*; RS: *Rhizoctonia solani*.

Additionally, pre-treatment of wheat seeds with *Delftia lacustris* strain NSC not only enhanced seed germination but also promoted wheat plantlet growth. Seeds treated with *Delftia lacustris* NSC exhibited significantly increased shoot length (*P = 0.033*) (**Figure 2C**) and root length (*P = 0.002*) (**Figure 2D**) relative to untreated seeds. Furthermore, it has significantly improved the root (*P = 0.020*, *P = 0.032*) and shoot length (*P = 0.031*, *P = 0.129*) of plantlets derived from wheat seeds pre-infected with *Rhizoctonia solani* and *Fusarium oxysporum*, respectively. Even after exposure to these pathogens, the average shoot and root lengths of plantlets treated with *Delftia lacustris* isolate NSC remained significantly higher (*P = 0.0297* and *P = 0.010*) compared to the untreated control, highlighting its biocontrol potential. Pre-treatment with these microbial strains not only.

The role of *Delftia lacustris* strain NSC in enhancing wheat seed germination under abiotic stress, such as high salinity, was also evaluated. Germination rates significantly decreased with increasing salt concentrations, up to 500mM NaCl (**Figure 3**). However, seeds pre-treated with *Delftia lacustris* strain NSC exhibited improved germination under high salinity conditions (**Figure 3A**). Specifically, seed germination increased by approximately 10.11-fold (*P = 0.0001*) at 1 M NaCl, with corresponding increases in alpha-amylase activity (0.412 IU at 1 M NaCl) (**Figure 3B**) compared to controls. Pre-treatment with it enhanced seed germination and promoted wheat plantlet growth. At the highest NaCl concentration (1 M), seed pretreatment exhibited a ∼4.02-fold increase in shoot length (*P = 0.0003*) (**Figure 3E**) and a ∼2.34-fold increase in root length (*P = 0.0001*) (**Figure 3F**) compared to untreated seeds under saline conditions (**Figure 3C, D**).

**Figure 3:**
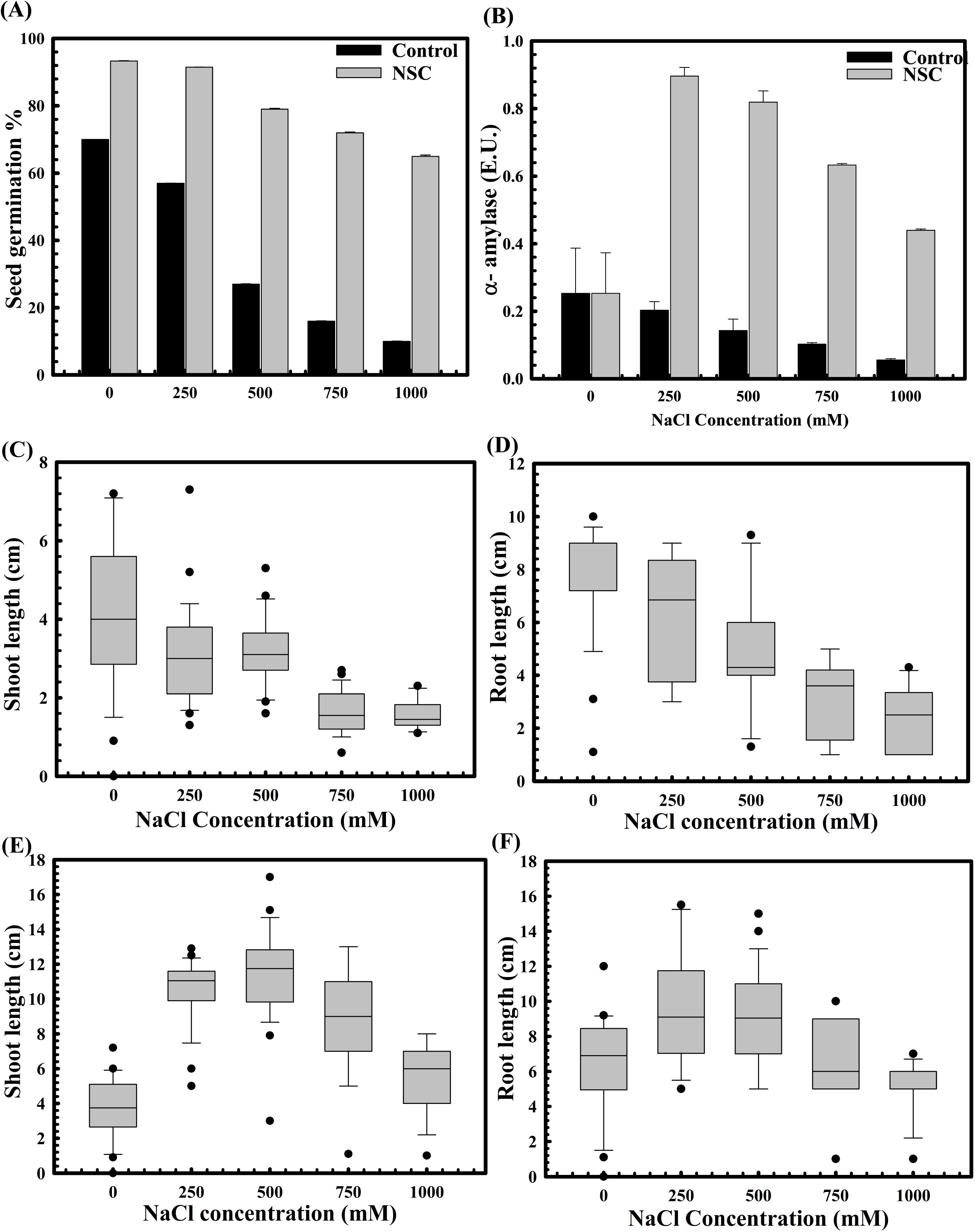
The impact of *Delftia lacustris* strain NSC pre-treatment on wheat seed germination in saline conditions. The seed germination assays were performed to compare the effect of *Delftia lacustris* strain NSC on seed germination under saline conditions (A). The alpha amylase activity of *Delftia lacustris* strain NSC pretreated wheat seeds under saline stress conditions (B). Shoot and root lengths of WC-306 seedlings were assessed under both untreated (C, D) and *Delftia lacustris* strain NSC pre-treated group (E, F) under NaCl-induced salinity stress. All assays were performed in triplicates.

### Assessment of *Delftia lacustris isolate* NSC biofertilizer properties under field conditions

The physiological parameters were assessed at the various growth stages of the wheat plant, also known as Feeks. The rhizosphere of wheat plants derived from seeds pre-treated with *Delftia lacustris* strain NSC exhibited substantial enhancements in various physiological parameters such as phosphate solubilization, nitrogen assimilation, the release of total sugar, and reducing sugar relative to untreated controls. Across multiple growth stages (Feeks 1, 2, 3, 6, 9, and 10.5), the concentrations of total sugars and reducing sugars in the rhizosphere were significantly elevated in the pre-treated plants, with total sugars increasing by approximately 2.1 to 3.20 times (**Figure 4A**), and reducing sugars by 0.10 to 1.09 times (**Figure 4B**). Likewise, extracellular alkaline phosphatase activity was markedly higher in the pre-treated plants, showing increases ranging from 3 to 6.66 times at different stages (**Figure 4C**). Additionally, pre-treatment resulted in enhanced nitrogen assimilation in the rhizosphere, as evidenced by a substantial increase in nitrate reductase activity. This activity was elevated by up to 9.05 times at Feeks 6, with consistent increases observed at other stages, ranging from 1.33 to 8 times higher than that in untreated seeds (**Figure 4D**). These findings indicate that the pre-treatment facilitated improved nutrient cycling, particularly in relation to sugars and phosphorus, and promoted greater nitrogen uptake, thereby supporting enhanced growth and development of the wheat plants. Wheat plants originated from seeds treated with *Delftia lacustris* strain NSC showed a significant increase in the number of tillers (*P=0.0042*), number of leaves per plant (*P=0.0002*), spike length (*P=0.0025*), number of spikes per plant (P=0.0033), number of spikelets per plant (*P=0.0001*), grain weight per 1000 grains (*P=0.0011*) and grain yield (*P=0.0001*) (**Table 2**). These findings suggested that *Delftia lacustris* strain NSC enhanced wheat productivity by promoting both vegetative growth (e.g., more tillers and leaves) and reproductive success (e.g., longer spikes, more spikelets, and higher grain weight), leading to higher overall grain yields. This could have important implications for sustainable agriculture, particularly as these microbial isolates may reduce the need for chemical fertilizers and enhance crop performance under varying environmental conditions.

**Figure 4:**
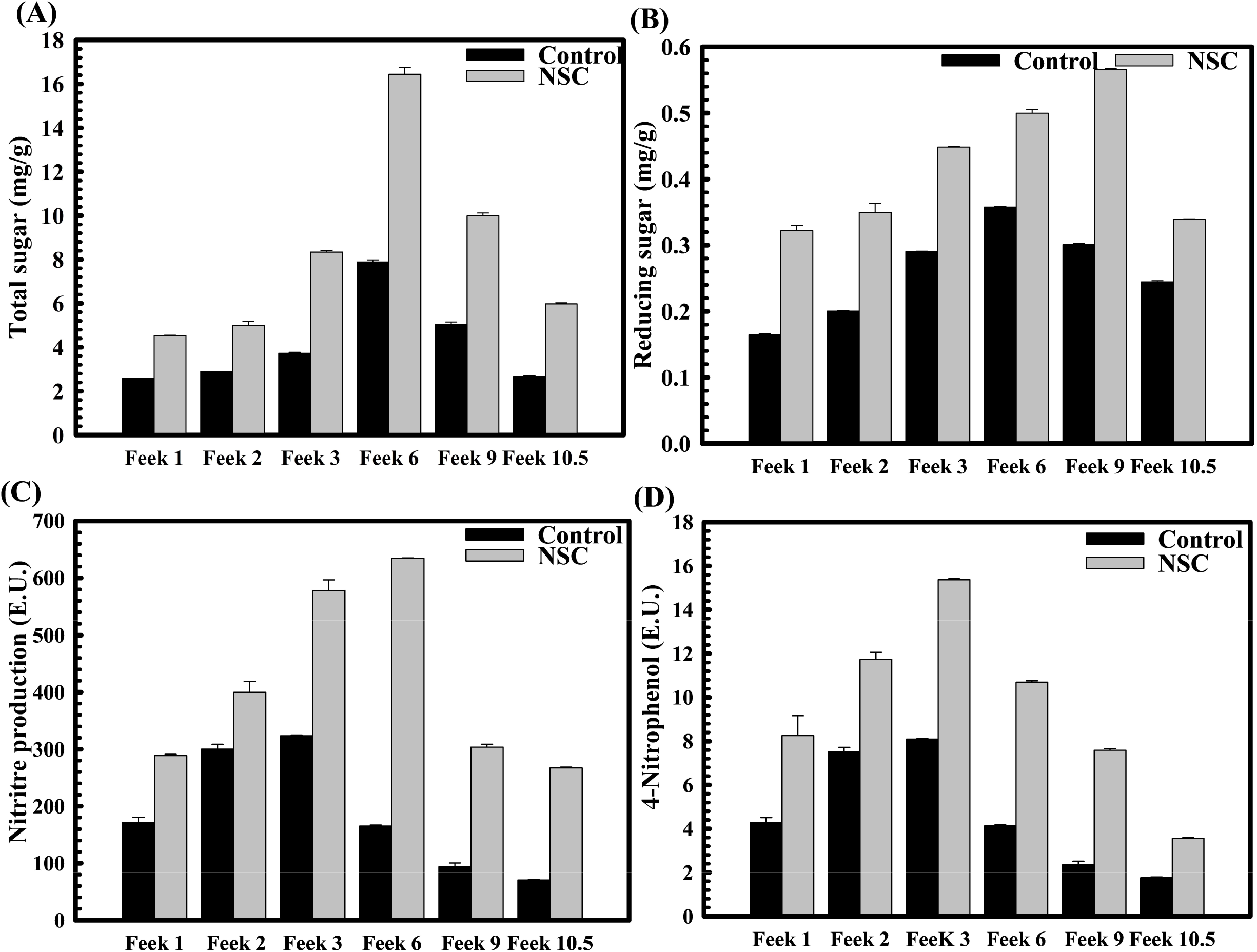
The impact of *Delftia lacustris* strain NSC pre-treatment on rhizosphere sugar content, phosphatase, and nitrate reduction activity across various wheat growth stages. Total sugar content was observed at different Feeks (1.0, 2.0, 3.0, 6.0, 9.0, and 10.5) in the presence and the absence of microbial inoculant NSC (A). Reducing sugar was estimated using DNS assay at different Feeks (1.0, 2.0, 3.0, 6.0, 9.0, and 10.5) was assessed in the presence and the absence of microbial inoculant NSC (B). Assessment of nitrate reductase activity at different Feeks (1.0, 2.0, 3.0, 6.0, 9.0, and 10.5) in the presence or absence of *Delftia lacustris* strain NSC (C). Assessment of alkaline phosphatase activity at different Feeks (1.0, 2.0, 3.0, 6.0, 9.0, 10.5) in the presence or absence of *Delftia lacustris* strain NSC (D). The experiment was carried out in triplicates. Experiments were carried out in triplicates Plotted values are the mean of triplicate readings and their observed standard deviation.

**Table 2:** Assessment of plant productivity features in *Delftia lacustris* strain NSC treated seeds in comparison to untreated. All the experiments were performed in triplicates. Values in the table represent the mean and S.D.

| Productivity phenotype | WC-306<br>Plants | WC-306 plants treated<br>with *Delftia lacustris* NSC |
| --- | --- | --- |
| Leaves per plant | 12.34±0.57 | 46 ±.05 |
| Number of tillers | 3.34±0.57 | 5.53±0.22 |
| Number of spike/plant | 26.3±1.309 | 49.9±1.2 |
| Spike length (cm) | 15.46±0.41 | 22.69±0.52 |
| Number of spikelets per<br>plant | 30.34±0.57 | 59.22±1.02 |
| Grain weight (g) | 30.23±0.37 | 58±0.57 |
| Grain yield (Kg/acre) | 3267 | 56328.5 |

## Discussion

An increase in wheat crop production is crucial to meet the rising food demands of the expanding global population (Hemathilake and Gunathilake, 2022). Traditionally, agrochemicals have been a primary strategy to improve agricultural yields and address food security concerns. However, the extensive and prolonged application of these chemicals has negative consequences for both the environment and human health. In particular, agrochemicals degrade soil fertility, which creates significant challenges for sustaining long-term crop productivity and securing food supplies (Kopittke et al., 2019). Additionally, the development of pesticide-resistant plant pathogens, which exhibit greater infectivity and wider host ranges, intensifies the difficulties faced by modern agriculture (Tudi et al., 2021). In response to these issues, sustainable agricultural practices have been proposed as a means to reduce the harmful effects of chemical inputs, promote healthy soil ecosystems, decrease pathogen pressures, and improve crop production (De Corato et al., 2024). Plants naturally host diverse communities of microorganisms on both their aerial and subterranean surfaces (Hao et al., 2024). These microbial communities promote plant growth by enhancing nutrient uptake, stimulating cell division, supporting reproductive processes, and providing defense against pathogen invasion (Kumar et al., 2022). Sustainable agriculture advocates for using these beneficial microbes to enhance crop productivity. Consequently, identifying and characterizing these beneficial microorganisms has become a focal point of global research as scientists aim to leverage their potential to improve agricultural sustainability and productivity.

*Delftia* species were identified from various habitats and are known to promote plant growth through enhanced nutrient assimilation and the extension of stress mitigation properties to the host (Braña et al., 2016). *Delftia* sp. are recognized as plant growth-promoting bacteria with a range of beneficial effects on plant growth and health (Han et al., 2005). Several strains of *Delftia* such as *Delftia lacustris*, *Delftia acidovorans, Delftia tsuruhatensis* can enhance plant development through mechanisms such as the production of phytohormones like indole-3-acetic acid (IAA), which promote root growth and improve nutrient uptake (Woźniak et al., 2019). Despite phenomenal plant growth and stress mitigation properties, *Delftia* species were not characterized for their role in wheat growth promotion. Hereby the current study aims to uncover candidate *Delftia* strains that can enhance wheat growth through mechanisms such as nutrient solubilization, hormone production, and pathogen suppression, thereby contributing to sustainable agricultural practices. Phylogenetic characterization of isolated train NSC demonstrated a high degree of homology with *Delftia lacustris*, thereby classifying it as *Delftia lacustris* strain NSC. In vitro, experiments confirmed the ability of *Delftia lacustris* strain NSC to solubilize phosphate, reduce nitrate, solubilize siderophore, and synthesize auxins and ammonia, which are key traits contributing to its biofertilizer potential. These properties are essential features of any biofertilizer strain and could enhance nutrient assimilation to improve host growth (Rao et al., 2020). The presence of these properties in *Delftia lacustris* strain NSC indicated it as a potential biofertilizer candidate to sustainably enhance wheat growth and production. Such properties were also identified in *Delftia* strains, enhancing plant growth and yield (Brambilla et al., 2022). *Delftia* species can solubilize phosphate, making this vital nutrient more accessible to plants (Elhaissoufi et al., 2022). Some strains also exhibit antagonistic activity against plant pathogens, contributing to disease suppression (Han et al., 2005). Likewise other *Delftia* strains, *Delftia lacustris* strain NSC also showcased prominent antifungal activity against phytoptahogens. Due to such multifaceted capabilities, *Delftia* sp. are being explored as potential bioinoculants to improve crop yield and sustainability in agriculture (Nikel, 2016). These studies reinforce the potential of *Delftia lacustris* strain NSC as a plant growth promoter and biocontrol agent. Most of the information is derived from experiments conducted on a variety of plant species. Therefore, it is crucial to investigate its physiological, genomic, and wheat plant-associated characteristics to fully comprehend its efficacy.

Accordingly, plant-growth-promoting and biocontrol characteristics of *Delftia lacustris* strain NSC were further validated through in vivo studies, specifically wheat seed germination. These experiments showed that the exposure of *Delftia lacustris* NSC effectively protected wheat seedlings from fungal infection, supporting its biocontrol potential. This suggests that the strain not only safeguards plants from disease but also fosters overall plant development. Furthermore, *Delftia lacustris* NSC improved seed germination and seedling growth under saline stress conditions, demonstrating its potential to mitigate the adverse effects of salinity on plant growth. These results highlight the versatile PGPR and biocontrol properties of *Delftia lacustris* strain NSC across a range of environmental stressors, positioning it as a promising candidate for sustainable agricultural practices. Pre-inoculation of wheat seeds with *Delftia lacustris* NSC resulted in a significant increase in plant growth-promoting features such as total sugar, reducing sugar, phosphate solubilization, and nitrate reduction. Microbial inoculation under field conditions also increased plant biomass and crop yield. All these results confirmed the successful translation of wheat growth promotion properties of *Delftia lacustris* strain NSC to the wheat host.

Biofertilizers must be thoroughly characterized for their stability in dynamic soil ecosystems (Márquez et al., 2023), plant growth-promoting capabilities (Sharma et al., 2024a), ability to establish stable colonization (Sharma et al., 2024a), and non-pathogenic behavior (Sharma et al., 2024a). Physiological stress response analysis of microbial isolate revealed its resilience under a range of environmental conditions, including varying pH (5–10), temperature (20– 55°C), drought, and salinity (>12% w/v), as well as exposure to arsenic. Their tolerance to these common soil stressors demonstrates their potential for consistent performance as plant growth promoters. Additionally, previous studies have reported the accumulation of antibiotics in soil, arising from sources such as farm animal manure, discarded pharmaceuticals, sludge, effluent water (Loh et al., 2018), or directly from soil microbes (Cycoń et al., 2019). Microbial isolate resisted several antibiotics, such as amoxycillin, bacitracin, cephalothin, vancomycin, ceftazidime, netilin, and ofloxacin. This resistance was further corroborated by the identification of antibiotic resistance genes through genome-wide analysis. These findings suggest that these isolates possess the ability to thrive in environments with elevated antibiotic concentrations, whether from human activities or microbial sources. Furthermore, the substrate utilization profile of both isolates highlights their metabolic versatility, as they can utilize a wide array of carbon sources, thereby enhancing their adaptability to environments with varied nutrient availability (Wang et al., 2019; Sharma et al., 2024a). Collectively, these characteristics confirm the isolates’ stability in complex soil ecosystems, a crucial property for effective biofertilizer functionality. Notably, the genome of *Delftia lacustris* NSC encodes several key hydrolytic enzymes, including proteases, chitinases, glucanases, and lipases, which are involved in the breakdown of complex organic materials. Additionally, it possesses the genetic potential to produce phytohormones like indole-3-acetic acid (IAA), solubilize essential nutrients such as phosphate and nitrogen, fix atmospheric nitrogen, and produce phenazines compounds with potent antifungal activity.

These genomic features position *Delftia lacustris* strain NSC as a strong candidate for use as a biofertilizer, as they contribute to various plant growth-promoting processes, including nutrient solubilization, hormone production, and disease suppression (Ubalde et al., 2012). Previous studies have demonstrated that strains of *Delftia* with similar characteristics are classified as plant growth-promoting rhizobacteria (PGPRs) due to their ability to enhance plant growth (Ubalde et al., 2012). The production of chitinolytic and proteolytic enzymes allows *Delftia* strains to degrade fungal cell walls, inhibiting fungal growth and providing biocontrol against plant pathogens (Nefzi et al., 2019). Moreover, phenazine production protects against fungal phytopathogens, as phenazines are well-documented for their antifungal properties (Karmegham et al., 2020).

The presence of genes encoding Type I and IV pili in the *Delftia lacustris* NSC genome further underscores its potential as a PGPR, as these pili play crucial roles in adhesion, autoaggregation, and biofilm formation, facilitating its interaction with plant roots in the rhizosphere (Zhang et al., 2024). This genome also suggests a strong interaction with the wheat rhizosphere, which is essential for promoting plant growth in this context. Despite the promising characteristics of *Delftia* species, concerns regarding their pathogenic potential have been raised, particularly for *Delftia lacustris* (Berg et al., 2005). However, the genome of *Delftia lacustris* strain NSC lacks genes typically associated with pathogenicity, thus confirming its safety for agricultural applications. These genomic and functional attributes indicate that *Delftia lacustris* NSC has significant potential as both a plant growth-promoting biofertilizer and a biocontrol agent, supporting sustainable agricultural practices. Utilizing this strain could offer a cost-effective and environmentally sustainable alternative to chemical fertilizers, enhancing plant nutrient availability while reducing the environmental impact associated with conventional fertilization practices.

## Author Contributions

NSC designed the study and experiments. NSC, PS, and RP wrote the manuscript. PS carried out the experiments. NSC, PS, and RP analyzed the data. All authors edited the manuscript and approved the final draft of the manuscript.

## Acknowledgment

The authors acknowledge CSIR-Institute of Genomics and Integrative Biology, New Delhi, India for DNA sequencing facility.

## Data Availability Statement

The 16S rRNA gene sequence datasets generated in this study were deposited at NCBI with SRA accession ID SUB15090314 and BioProject ID: PRJNA1223623 (https://www.ncbi.nlm.nih.gov/sra/PRJNA1223623).

