## Supplementary tables and Figures for "Assessing wheat growth promotion potential of *Delftia lacustris* strain NSC through genomic and physiological characterization"

**Running Title:** *Delftia lacustris as* wheat biofertilizer.

**Number of Words: 7542**
**Number of Figures: 4**
**Number of Tables: 2**

**Supplementary Table S1:** Substrate utilization profile of *Delftia lacustris* strain NSC with other phylogenetic-related *Delftia* species

| Sr. No. | Substrate | *Delftia lacustris* NSC | *Delftia lacustris* LzhVag01 | *Delftia tsuruhatensis* CM13 | *Delftia tsuruhatensis* NBRC 16741 | *Delftia acidovorans* |
|---|---|---|---|---|---|---|
| 1 | Lactose | + | + | ND | ND | ND |
| 2 | Xylose | + | ND | ND | ND | ND |
| 3 | Maltose | + | ND | ND | ND | + |
| 4 | Fructose | ND | + | + | ND | ND |
| 5 | Dextrose | + | + | + | ND | + |
| 7 | Raffinose | + | ND | ND | ND | ND |
| 8 | Trehalose | + | + | ND | ND | ND |
| 9 | Melibiose | ND | ND | ND | ND | ND |
| 10 | Sucrose | + | + | ND | ND | ND |
| 11 | L-Arabinose | ND | ND | ND | ND | + |
| 12 | Mannose | ND | ND | + | ND | + |
| 13 | Inulin | + | + | ND | ND | ND |
| 14 | Glycerol | ND | + | ND | ND | ND |
| 15 | Dulcitol | + | ND | ND | ND | ND |
| 16 | Mannitol | + | + | ND | ND | + |
| 17 | Adonitol | + | ND | ND | ND | ND |
| 18 | Rhamnose | + | ND | ND | ND | ND |
| 19 | Xylitol | + | ND | + | ND | ND |
| 20 | ONPG | + | + | ND | ND | ND |
| 21 | Esculin hydrolysis | ND | ND | ND | + | ND |
| 22 | D-Arabinose | ND | ND | ND | ND | ND |
| 23 | Citrate utilization | ND | ND | ND | + | ND |
| 24 | Malonate utilisation | ND | ND | ND | ND | ND |
| 25 | Sorbose | + | + | ND | ND | + |

Here ND: Not defined in the literature.

**Supplementary Table S2:** Comparative analysis of antibiotic susceptibility profile of *Delftia lacustris* NSC with other phylogenetic similar *Delftia* strains.

| Sr. No. | Antibiotic | ***Delftia lacustris* NSC** | ***Delftia lacustris* LzhVag01** | ***Delftia tsuruhatensis* CM13** | ***Delftia tsuruhatensis* NBRC 16741** | ***Delftia acidovorans*** |
|---|---|---|---|---|---|---|
| 1 | Amikacin | ND | ND | + | + | ND |
| 2 | Amoxicillin | ND | ND | ND | + | ND |
| 3 | Bacitracin | + | + | ND | ND | ND |
| 4 | Cephalothin | ND | ND | ND | ND | ND |
| 5 | Erythromycin | ND | + | ND | + | ND |
| 6 | Novobiocin | + | + | ND | ND | ND |
| 7 | Oxytetracyclin | + | ND | ND | ND | |
| 8 | Vancomycin | + | + | ND | ND | ND |
| 9 | Ceflnaxone | ND | ND | + | ND | ND |
| 10 | Ceftazidime | + | + | ND | ND | + |
| 11 | Cefotaxime | ND | ND | + | ND | ND |
| 12 | Lincomycin | ND | + | ND | ND | ND |
| 13 | Netillin | ND | ND | + | ND | ND |
| 14 | Ofloxacin | + | ND | ND | ND | ND |

Here, ND is not defined in the literature.

**Supplementary Table S3:** A Comparison of the growth parameters of *Delftia lacustris* NSC with other *Delftia* strains

| Sr no. | Microbes | pH | Temperature (˚C) | Minimum Inhibitory Concentration | | | | | | |
|---|---|---|---|---|---|---|---|---|---|---|
| | | | | **NaCl** | **KCl** | **LiCl** | **As(III)** | **As(V)** | **$CdCl_2$** | **$H_2O_2$** |
| 1 | ***Delftia lacustris* NSC** | 5-10 | 20-55 | 1750mM | 2000 mM | 1750 mM | 1500 PPM | 1000 PPM | 6mM | 12.5mM |
| 2 | ***Delftia lacustris* LzhVag01** | 6-9 | 15-30°C | 102.7mM | ND | ND mM | ND | 1000 PPM | ND | ND |
| 3 | ***Delftia tsuruhatensis* CM13** | 5 to 9 | 20 to 40 | 900mM | ND | ND | ND | ND | ND | ND |
| 4 | ***Delftia tsuruhatensis* NBRC 16741** | 5-10 | 20–45 | 1000mM | ND | ND | ND | ND | ND | ND |
| 5 | ***Delftia acidovorans*** | 5-10 | 20–40 | 530mM | ND | ND | ND | ND | ND | ND |

Here ND: Not defined in the literature.

**Supplementary Table S4**: Average nucleotide identity (ANI) of *Delftia lacustris* NSC with other *Delftia* species.

| | **Microbial isolate NSC** | ***Delftia acidovorans*** | ***Delftia lacustris*** | ***Delftia* sp hk171** | ***Delftia tsuruhatensis* NBRC 16741** | ***Delftia tsuruhatensis T7*** |
|---|---|---|---|---|---|---|
| **Microbial isolate NSC** | * | 94.54 | **98.08** | 94.65 | 98.05 | 93.16 |
| ***Delftia acidovorans*** | 94.48 | * | 94.59 | 97.4 | 81.45 | 89.92 |
| ***Delftia lacustris*** | **98.28** | 94.34 | * | 94.39 | 84.85 | 92.62 |
| ***Delftia* sp hk171** | 94.42 | 97.31 | 94.53 | * | 80.95 | 89.51 |
| ***Delftia tsuruhatensis* NBRC 16741** | 97.65 | 95.05 | 96.45 | 93.93 | * | 95.79 |
| ***Delftia tsuruhatensis* T7** | 97.68 | 93.87 | 96.76 | 93.73 | 88.31 | * |

**Supplementary Table S5:** Tetra correlation among *Delftia lacustris NSC* and other *Delftia* species by a wide distribution of Z-score.

| **Organism** | **Z- Score** |
|---|---|
| *Delftia lacustris* LMG 24775 | 0.99989 |
| *Delftia tsuruhatensis* CM13 | 0.99983 |
| *Delftia tsuruhatensis* NBRC 16741 | 0.99976 |
| *Delftia acidovorans* CCUG 15835 | 0.99976 |
| *Delftia acidovorans* CCUG 274B | 0.99975 |
| *Delftia* sp. 670 | 0.99964 |
| *Delftia* sp. Cs1-4 | 0.99959 |
| *Delftia* sp. JD2 | 0.99957 |
| *Delftia tsuruhatensis* 391 | 0.99955 |
| *Delftia acidovorans* SPH-1 | 0.99955 |
| *Delftia acidovorans* FDAARGOS_997 | 0.9995 |
| *Delftia acidovorans* 2167 | 0.99945 |
| *Delftia acidovorans* NBRC 14950 | 0.99933 |
| *Delftia tsuruhatensis* MTQ3 | 0.99925 |
| *Delftia* sp. RIT313 | 0.99913 |
| *Comamonas terrae* NBRC 106524 | 0.98431 |
| *Comamonas terrae* TISTR 1906 | 0.9843 |
| *Comamonas phosphati* CGMCC 1.12294 | 0.98044 |
| *Acidovorax* sp. CF316 | 0.97175 |
| *Xenophilus arseniciresisten*s YW8 | 0.96631 |
| *Acidovorax* sp. Root217 | 0.96576 |
| *Acidovorax* sp. Root219 | 0.96546 |
| *Paenacidovorax monticola* KACC 19171 | 0.9651 |
| *Pseudorhodoferax soli* DSM 21634 | 0.96355 |
| [*Acidovorax*] ebreus TPSY | 0.96319 |
| *Pseudorhodoferax* sp. Leaf267 | 0.96281 |
| *Diaphorobacter nitroreducens* DSM 15985 | 0.96259 |
| *Alicycliphilus denitrificans* K601 | 0.96209 |
| *Acidovorax* sp. JS42 | 0.96199 |
| *Alicycliphilus* sp. B1 | 0.96192 |
| *Diaphorobacter* sp. J5-51 | 0.96168 |
| *Pseudorhodoferax* sp. Leaf274 | 0.96126 |
| *Alicycliphilus denitrificans* BC | 0.96087 |
| *Pseudorhodoferax* sp. Leaf265 | 0.96075 |
| *Pseudorhodoferax aquiterrae* KCTC 23314 | 0.9592 |
| *Acidovorax* sp. MR-S7 | 0.95598 |

| | |
|---|---|
| *Comamonas aquatica* NBRC 14918 | 0.95559 |
| *Comamonas aquatica* DA1877 | 0.95538 |
| *Comamonas granuli* NBRC 101663 | 0.95212 |
| *Simplicispira lacusdiani* CPCC 100842 | 0.95101 |
| *Variovorax terrae* CYS-02 | 0.94929 |
| *Comamonas guangdongensis* CCTCC AB2011133 | 0.94658 |
| *Pulveribacter suum* SC2-7 | 0.94495 |
| *Paracidovorax wautersii* DSM 27981 | 0.9425 |
| *Paracidovorax anthurii* CFPB 3232 | 0.94081 |
| *Paracidovorax konjaci* DSM 7481 | 0.93916 |
| *Comamonas testosteroni* TK102 | 0.93781 |
| *Diaphorobacter limosus* Y-1 | 0.9371 |
| *Simplicispira sedimenti* W1-6 | 0.93708 |
| *Acidovorax soli* DSM 25157 | 0.93689 |
| *Aquabacterium soli* SJQ9 | 0.93507 |
| *Comamonas testosteroni* KF-1 | 0.93505 |
| *Acidovorax* sp. Leaf191 | 0.93494 |
| *Acidovorax* sp. Leaf76 | 0.93492 |
| *Acidovorax* sp. Leaf84 | 0.93414 |
| *Curvibacter* sp. PAE-UM | 0.93308 |
| *Ramlibacter rhizophilus* CCTCC AB2015357 | 0.93306 |
| *Hydrogenophaga borbori* LA-38 | 0.93269 |
| *Comamonas testosteroni* JL40 | 0.93219 |
| *Paracidovorax cattleyae* DSM 17101 | 0.93216 |
| *Paenacidovorax caeni* R-24608 | 0.93196 |
| *Paracidovorax oryzae* ATCC 19882 | 0.93158 |
| *Paenacidovorax caeni* R-24608 | 0.93154 |
| *Paracidovorax avenae* ATCC 19860 | 0.93071 |
| *Comamonas endophytica* 5MLIR | 0.9303 |
| *Acidovorax delafieldii* 2AN | 0.93007 |
| *Paracidovorax valerianellae* DSM 16619 | 0.92985 |
| *Comamonas terrigena* NBRC 13299 | 0.92962 |
| *Comamonas terrigena* NBRC 13299 | 0.92908 |
| *Acidovorax* sp. Leaf78 | 0.92891 |
| *Comamonas terrigena* NCTC1937 | 0.92888 |
| *Comamonas testosteroni* WDL7 | 0.92886 |
| *Variovorax soli* NBRC 106424 | 0.9287 |
| *Melaminivora suipulveris* SC2-9 | 0.92835 |
| *Comamonas thiooxydans* DS1 | 0.92826 |
| *Paracidovorax citrulli* AAC00-1 | 0.92712 |

| | |
|---|---|
| *Bordetella hinzii* 5132 | 0.92707 |
| *Comamonas resistens* ZM22 | 0.92684 |
| *Comamonas thiooxydans* JL14 | 0.92648 |
| *Comamonas antarctica* 16-35-5 | 0.92631 |
| *Paracidovorax citrulli* ICMP 7500 | 0.92627 |
| *Comamonas thiooxydans* DF1 | 0.92603 |
| *Bordetella hinzii* 1277 | 0.92599 |
| *Comamonas thiooxydans* JC9 | 0.92593 |
| *Paracidovorax citrulli* DSM 17060 | 0.92586 |
| *Bordetella hinzii* NCTC13199 | 0.92582 |
| *Comamonas* sp. E6 | 0.9258 |
| *Comamonas thiooxydans* DF2 | 0.92573 |
| *Comamonas thiooxydans* JC13 | 0.92573 |
| *Bordetella hinzii* F582 | 0.9257 |
| *Comamonas thiooxydans* DSM 17888 | 0.9257 |
| *Bordetella hinzii* L60 | 0.92569 |
| *Bordetella hinzii* LMG 13501 | 0.92568 |
| *Comamonas thiooxydans* DSM 17888 | 0.92558 |
| *Comamonas thiooxydans* DSM 17888 | 0.92558 |
| *Bordetella hinzii* OH87 BAL007II | 0.9255 |
| *Comamonas thiooxydans* JC12 | 0.92546 |
| *Comamonas thiooxydans* JC8 | 0.92536 |
| *Acidovorax* sp. Root275 | 0.92513 |

77
